# Linear versus Nonlinear Muscle Networks: A Case Study to Decode Hidden Synergistic Patterns During Dynamic Lower-limb Tasks

**DOI:** 10.1101/2023.01.15.524160

**Authors:** Rory O’Keeffe, Vaibhavi Rathod, Seyed Yahya Shirazi, Sarmad Mehrdad, Alexis Edwards, Smita Rao, S. Farokh Atashzar

## Abstract

This paper, for the first time, compares the behaviors of nonlinear versus linear muscle networks in decoding hidden peripheral synergistic neural patterns during dynamic functional tasks. In this paper, we report a case study during which one healthy subject conducts a series of four lower limb repetitive tasks. Specifically, the paper focuses on tasks that involve the right knee joint, including walking, sit-to-stand, stepping, and drop-jump. Twelve muscles were recorded using the Delsys Trigno system. The linear muscle network was generated using coherence analysis, and the nonlinear network was generated using Spearman’s correlation. The results show that the degree, clustering coefficient, and global efficiency of the muscle network have the highest value among tasks in the linear domain for the walking task, while a low linear synergistic network behavior for the sit-to-stand is observed. On the other hand, the results show that the nonlinear functional muscle network decodes high connectivity (degree) and clustering coefficient and efficiency for the sit-to-stand when compared with other tasks. We have also developed a two-dimensional functional connectivity plane composed of linear and nonlinear features and shown that it can span the lower-limb dynamic task space. The results of this paper for the first time highlight the importance of observing both linear and nonlinear connectivity patterns, especially for complex dynamic tasks. It should also be noted that through a simultaneous EEG recording (using BrainVision System), we have shown that, indeed, cortical activity may indirectly explain highly-connected nonlinear muscle network for the sit-to-stand task, highlighting the importance of nonlinear muscle network as a neurophysiological window of observation beyond the periphery.

## I. Introduction

Surface electromyography (sEMG) has been utilized in the literature to detect various muscle activation and formulate biomarkers of neuromuscular health or disease progression [1]. In the literature, classic muscle synergies have been typically utilized as a low-dimensional activation representation that suggests how the nervous system may combine a series of patterns to produce a complex movement [2]. Generalizing on this concept and focusing on a full-spectrum pattern of synergistic information sharing at the periphery, functional muscle connectivity has attracted a great deal of interest in the last decade [3], [4], [5], [6], [7] initially for healthy subjects and recently for patients with neural damages. We have recently shown that in stroke subjects [7], muscle networks can robustly detect motor improvement during the course of rehabilitation when conventional spectrotemporal features of sEMG fail to provide the needed robustness and accuracy. Beyond peripheral functionality, the muscle network decodes the distribution of the descending neural drive rooted in the central nervous system, which would result in hidden synergistic patterns during functional tasks. Despite the benefit, most of the existing literature utilizes linear spectral coherence analysis to decode functional connectivity [3], [4], [5], [6]. The use of magnitude coherence for producing muscle networks is motivated by the wide use of such measures in Brain-Computer Interfaces. However, it should be highlighted that due to the complex characteristics of volume conductors for muscle networks, different from relatively homogenous and small-size volume conductors in the brain, the synergistic coupling between various muscles may or may not follow a linear pattern. The authors have recently and for the first time, proposed the concept of a nonlinear muscle network using information theory for upper-limb tasks of stroke subjects [7]. Unlike linear connectivity metrics, such as Pearson’s correlation or coherence, a nonlinear technique (such as Spearman’s correlation or Mutual Information) can detect complex “information synchronization” in the distributed peripheral nervous activities. Motivated by the above-mentioned note, in this paper, we investigate the importance of linear versus nonlinear networks in detecting the hidden synergistic functional patterns in network scans. For this, the study includes data from one healthy subject who conducted a series of functional lower limb tasks, including walking, repeated sit-to-stand, repeated stepping, and repeated drop jump. The tasks were chosen because of their high relevance to daily living, recreational activity, current rehabilitation research, and clinical settings [8], [9], [10], [11]. Surface electromyography (sEMG) was collected from 12 muscles, while in parallel, 64-channel electroencephalography (EEG) was collected from the brain. In this paper, we utilize median magnitude coherence from 10Hz to 50Hz to produce the linear muscle network, while Spearman’s correlation was used to generate the nonlinear network scans. Spearman’s correlation models monotonic but nonlinear synchronicity in the activations of muscles which may not be detected by the coherence analysis. This paper is specifically interested to understand and uncover the potential of the nonlinear muscle network generated using Spearman’s correlation to measure variation in motor control over four lower-limb dynamic tasks. The collection of EEG is to evaluate how the cortical activation map relates to the linear versus nonlinear network behaviors. In order to analyze network behavior, in this paper, functional integration and segregation of the network scans are analyzed, which can evaluate changes in the collective neural drive, and are quantified using global efficiency (GE), degree, and weighted clustering coefficient (WCC), respectively (definitions of GE, degree, and WCC can be found in [12], [13]).

It should be noted that in this paper, we focus on the knee joint, and the central tasks (among four tested tasks) are sit-to-stand and walking. This choice of sit-to-stand is supported by previous works, which have shown such a task is a useful test of the health of the lower extremity [14], [15] when knee extension plays an important role [16], [17]. Due to the essential nature of these tasks in everyday life, metrics relating to sit-to-stand and walking are analyzed in clinical settings [18], [19], [20] and hence results from this work could inform rehabilitation programs.

The rest of this paper is organized as follows. In Section II, the implemented method is explained. Results and discussions are provided in Section III. And concluding remarks are provided in Section IV.

## II. Methods

### A. Experimental Procedure

One asymptomatic subject performed four lower limb dynamic tasks while wireless surface electromyography (sEMG) and 64-channel electroencephalography (EEG) were recorded. The subject performed the following tasks: (i) bilateral walking for 3 minutes, (ii) bilateral sit-to-stand for 45s, (iii) repetitions of unilateral step-up for 3 minutes, and (iv) ten repetitions of unilateral drop and jump. The bilateral tasks (walking and sit-to-stand) were conducted at a natural pace, and the unilateral tasks were performed on the right side. The sEMG signals from twelve sensors were recorded using the wireless Trigno sEMG system (Delsys Inc., Natick, MA), with a sampling frequency of 2148 Hz and an embedded 10 Hz high-pass filter.

Twelve bipolar Trigno Avanti sensors were utilized on both sides for (i) Gastrocnemius (GA), (ii) Tibialis Anterior (TA), (iii) Semitendinosus (ST), (iv) Rectus Femoris (RF), (v) Gluteus Maximus (GX) and (vi) Gluteus Medius (GD). The skin was wiped vigorously before placing the sensors.

sEMG sensors were oriented in line with the direction of the muscles. The knee joint angle on the right side was simultaneously monitored with a goniometer. EEG signals were wirelessly recorded from 64 electrodes assembled on an active-electrode cap (ActiCap, Brain Products GmbH, Germany) using a 10-10 system. Signals were processed using MATLAB R2020b (MathWorks Inc., Natick, MA) after the recording.

### B. Data Processing

A zero-phase high-pass filter was applied to the EEG signals at 0.1 Hz. A zero-phase low-pass filter was applied to both the EMG and EEG signals at 50 Hz, such that the resultant sEMG and EEG signals were in the 10-50 and 0.1-50 Hz ranges, respectively. Outlier channels were rejected if they exceeded a threshold value of kurtosis, and spherical interpolation was performed to replace the rejected channel data [21], [22]. Independent component analysis was used to remove artifacts from the EEG. The independent components were ranked by likelihood of being true brain signals with the ICLabel algorithm, and the top 50% most likely components were maintained [23]. The tasks were divided into epochs for calculating the intermuscular connectivity and cortical activity. The peaks of the right knee extension angle were recorded on the goniometer and delineated the epochs for walking, sit-to-stand, and unilateral stepping. The start and end time of the *i*^*th*^ epoch were the respective times at which the *i*^*th*^ and (*i* + 1)^*th*^ peaks occurred. The peaks of the moving average (*n* = 1000 samples) of rectified activation for the GD R were used to epoch the drop-jump task. The start and end of the time of the *i*^*th*^ epoch were identified as when the moving average increased beyond and decreased below 10% of the *i*^*th*^ peak. The sEMG and EEG signals were synchronized, and identical epoch times divided the tasks for analyzing cortical activity and intermuscular connectivity.

### C. Linear and nonlinear Muscle Networks

Linear and nonlinear muscle networks were constructed for each epoch of the dynamic tasks. Each node in the network represents a muscle, and the connection between nodes represents the synchrony between muscles. Intermuscular connectivity was computed with (i) magnitude squared coherence *C*_*xy*_ and (ii) Spearman power correlation *ρ*_*xy*_, which quantified linear and nonlinear synchrony, respectively.

To construct the linear muscle networks, *C*_*xy*_ between two signals *x*(*t*) and *y*(*t*) was computed as:

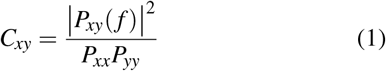

where *P*_*xx*_ and *P*_*yy*_ are the power spectral densities (PSDs) and *P*_*xy*_ is the cross power spectral density (CPSD) [neck paper]. To compute the coherence, Welch’s overlapped averaged periodogram method [24] was utilized with a Hamming window of 512 samples (238 ms) and 50% overlap. The median coherence component in the 20-50 Hz was selected for each sensor pair. Using this median coherence value, an adjacency matrix that represents the linear muscle network was constructed for each epoch.

In order to construct the nonlinear muscle networks, *ρ*_*xy*_ between two signals *x*(*t*) and *y*(*t*) was calculated. The power time series were obtained by computing *x*^2^(*t*), *y*^2^(*t*). Subsequently, each power time series with *n* samples was transformed with a rank operation such that the values ascended from 1 to *n*. After rank transforming *x*^2^(*t*) to *p*_*x*_(*t*) and *y*^2^(*t*) to *p*_*y*_(*t*), *ρ*_*xy*_ is computed according to:

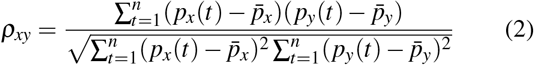

The magnitude *ρ*_*xy*_ was computed between each muscle pair across the epoch duration. Hence, the monotonic negative correlation between *x*^2^(*t*) and *y*^2^(*t*) of − *ρ*_*xy*_ is viewed as possessing equivalent non-parametric connectivity to the monotonic positive correlation of *ρ*_*xy*_. Using |*ρ*_*xy*_|, an adjacency matrix that represents the nonlinear muscle network was constructed for each epoch.

With respect to linear and nonlinear methods, the median adjacency matrix across epochs was used to quantify the intermuscular connectivity during the task. The twelve-muscle and the right-sided six-muscle adjacency matrices were considered for the bilateral and unilateral tasks, respectively (Figs. 1b, 2, 3). The degree, mean degree, mean weighted clustering coefficient, and global efficiency values of the linear and nonlinear muscle networks were quantified based on the respective adjacency matrices (Figs. 4, 5). The t-test was used to test the null hypothesis at the 0.05 significance level. The Bonferroni correction was applied to correct for multiple comparisons. The two-dimensional functional connectivity subspace was constructed using *C*_*xy*_ and |*ρ*_*xy*_| on the horizontal and vertical axes, respectively (Fig. 6). Each element of the adjacency matrices during a particular task is represented by a dot within the scatter plot.

**Fig. 1.**
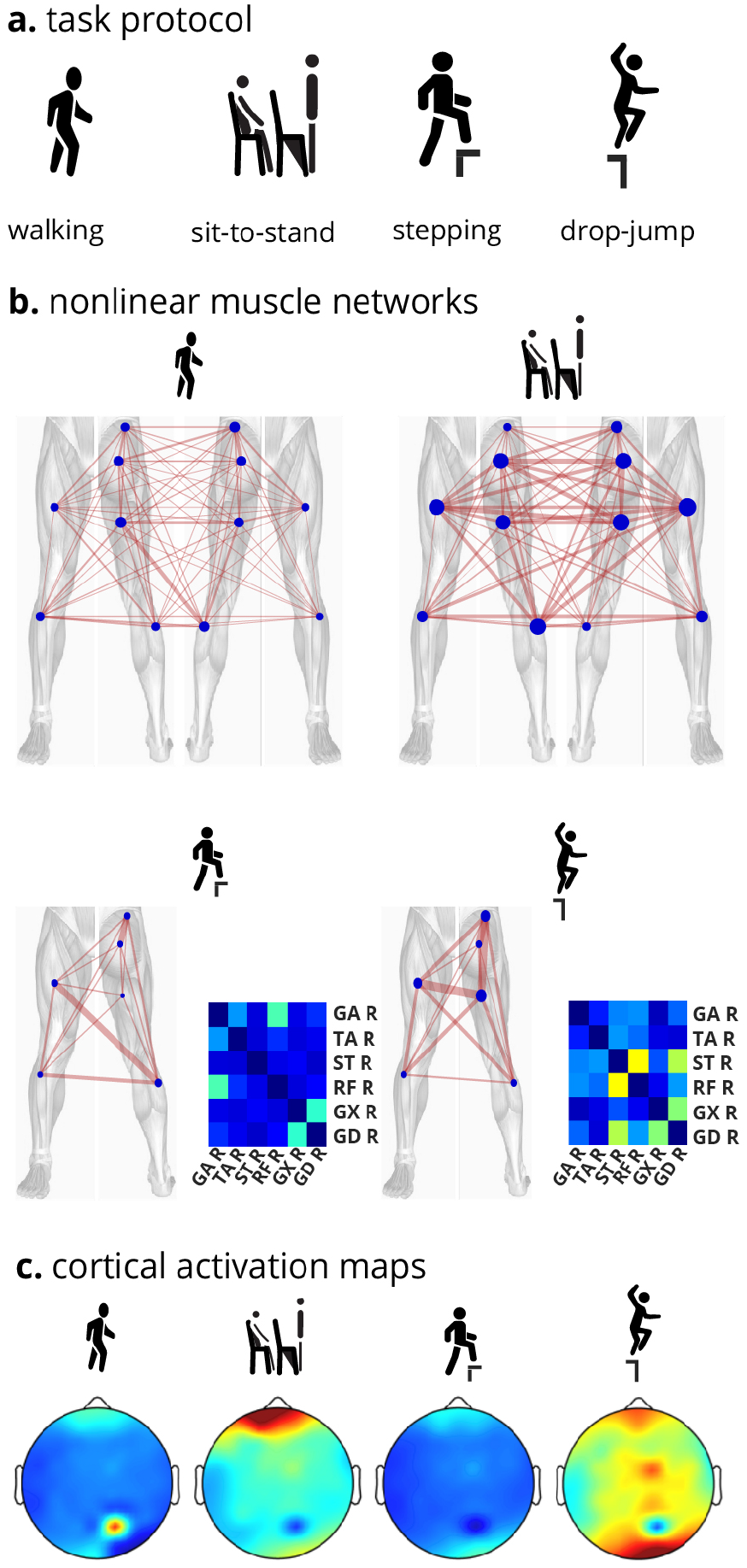
Study Overview: **a**. One asymptomatic subject performed four lower limb dynamic tasks while wireless surface electromyography (sEMG) and 64-channel electroencephalography (EEG) were recorded. **b**. sEMG sensors were placed bilaterally on the following muscles: (i) Gastrocnemius (GA), (ii) Tibialis Anterior (TA), (iii) Semitendinosus (ST), (iv) Rectus Femoris (RF), (v) Gluteus Maximus (GX) and (vi) Gluteus Medius (GD). Muscle networks (in this figure nonlinear muscle network is shown) and the corresponding adjacency matrices were constructed for each task using Spearman correlation (more details later in the paper). **c**. Cortical activations were quantified across the 64-channel. Pre-frontal and motor cortex activations appeared highest for sit-to-stand and drop-jump, respectively.

**Fig. 2.**
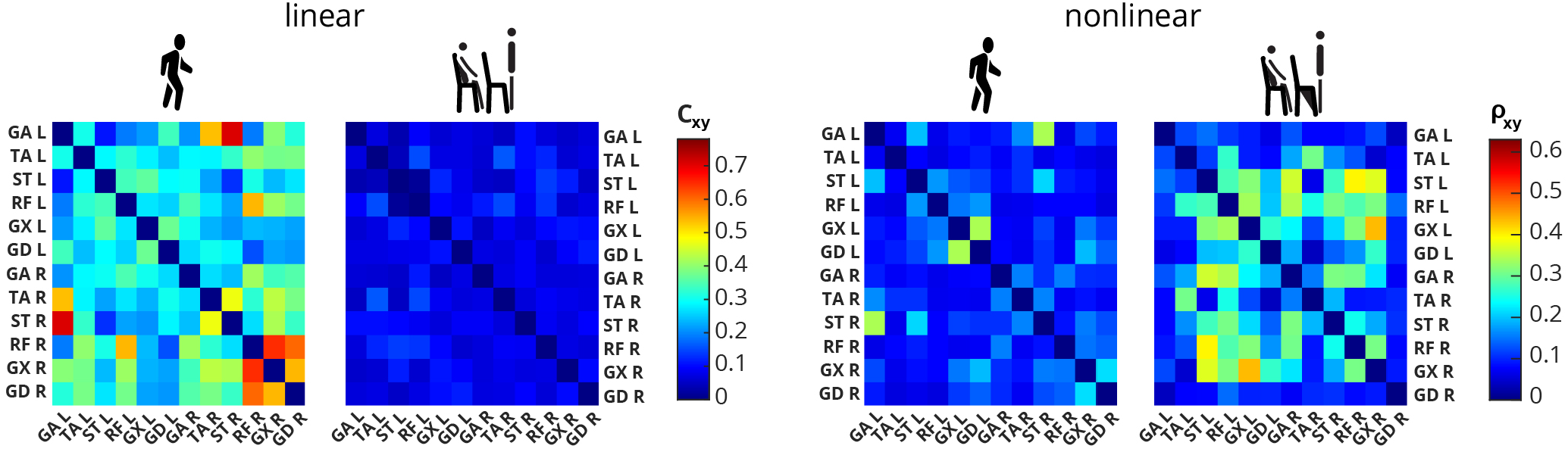
Twelve-muscle linear and nonlinear muscle networks were compared for the bilateral tasks (sit-to-stand and walking). The linear muscle network calculated using coherence appears to have higher connectivity trends for walking compared to sit-to-stand. On the other hand, the nonlinear muscle network calculated using Spearman’s power correlation showed trends of higher connectivity for sit-to-stand.

**Fig. 3.**
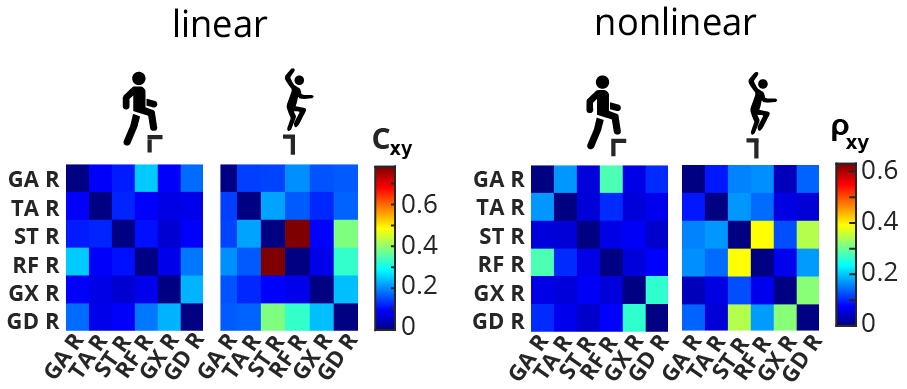
Six-muscle linear and nonlinear muscle networks were compared for the unilateral tasks (stepping and drop-jump). Both the linear and nonlinear muscle networks appeared to show higher connectivity for drop-jump.

**Fig. 4.**
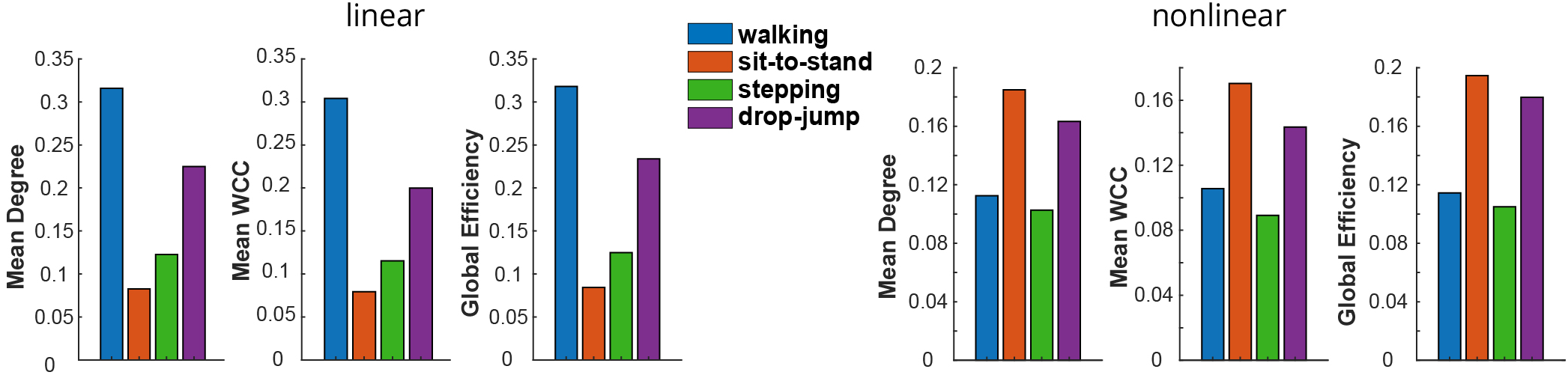
Linear muscle networks show walking while nonlinear muscle networks show sit-to-stand as having the highest functional integration out of the four tasks. Graph theory metrics (mean degree, mean WCC, and global efficiency) quantified the muscle networks’ functional integration and segregation. Drop-jump showed the second-highest functional integration for both linear and nonlinear methods.

**Fig. 5.**
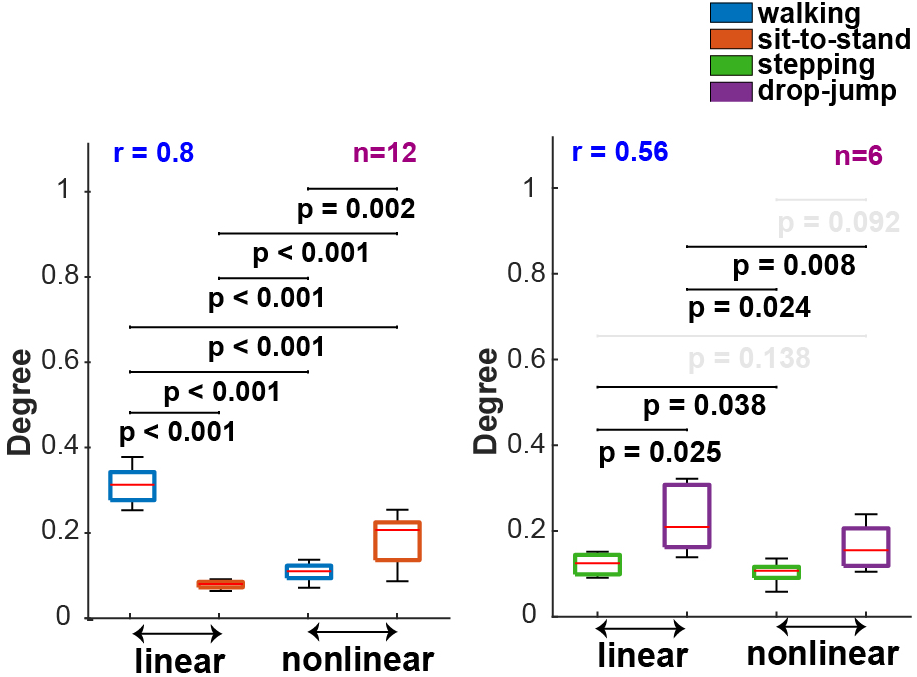
The node degree was higher for walking in the linear vs. nonlinear network comparison (t-test: *p <* 0.001). Meanwhile, the degree was higher for sit-to-stand when comparing linear and nonlinear networks (t-test: *p <* 0.001).

**Fig. 6.**
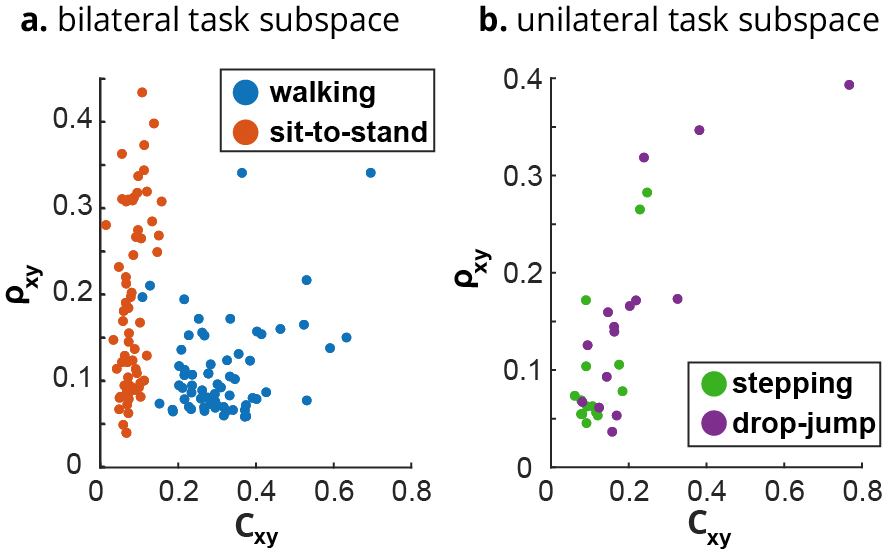
The two-dimensional connectivity subspace was constructed using coherence on the horizontal axis and Spearman correlation on the vertical axis. Each dot in the scatter plot represents an element of the corresponding adjacency matrix. The 2-D bilateral task subspace could separate walking and sit-to-stand for nearly all of the network edges.

### D. Cortical Activity

Cortical activity was quantified for each EEG channel as the root mean square (RMS) across the epoch. The median across epochs was used to construct the cortical activity heat maps (Fig. 1c). The motor cortex activations for each task were quantified with two metrics, (i) RMS and (ii) maximum absolute value *max*(*abs*) (Fig. 7). Each metric was computed for a 5 × 5 grid (indicated by a dashed rectangle on each mini-map) of electrodes centered at the Cz electrode, and the bar represents the median across the grid.

**Fig. 7.**
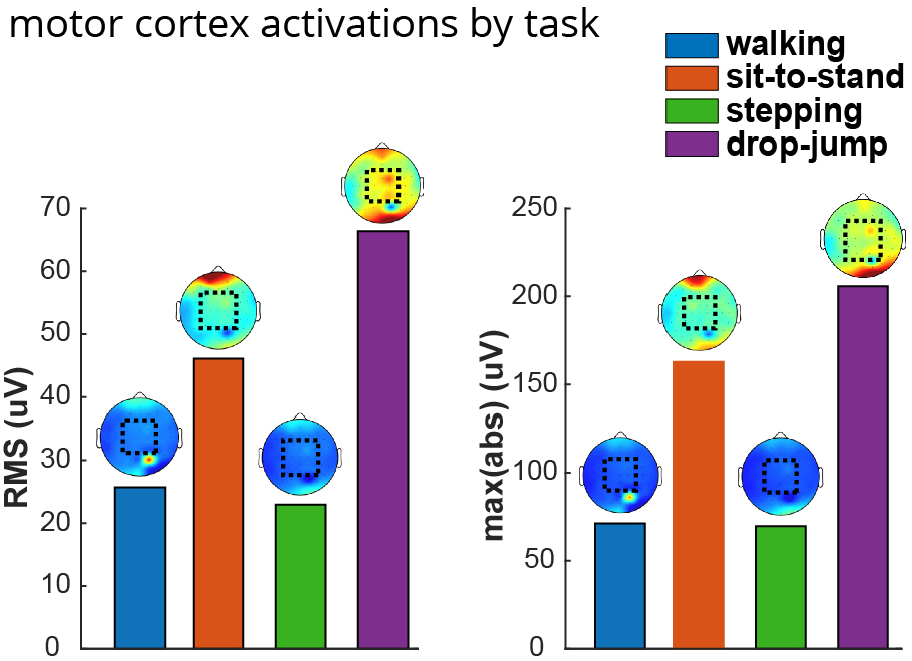
Motor cortex activations were highest The motor cortex activations for each task were quantified with two metrics, (i) RMS and (ii) max(abs). Each metric was computed for a 5 × 5 grid of electrodes centered at the Cz electrode, and the bar represents the median across the grid. The motor cortex activation during the sit-to-stand task was higher than for walking, as was the case for the nonlinear muscle network.

## III. Results

Fig. 2 represents the functional muscle network for bilateral tasks decoded using both linear and nonlinear connectivity methods. Fig. 3 represents the small unilateral network for stepping and drop-jump tasks. The small network excludes cross-sagittal connectivity. The results highlight that muscle network scans of the lower limb encode different synergistic patterns when comparing the tasks.

The most important observation from the results is that linear and nonlinear muscle networks showed opposing patterns of functional integration for the bilateral tasks (Figs. 2, 4, 5, 6) while agreeing on the unilateral tasks. In this regard, it should be noted that the linear, coherence-based network showed a stronger and more efficient (considering the global efficiency) network and higher clustering coefficient for the walking task (degree, *t-test: p <* 0.001, Fig. 5). However, the nonlinear network showed increased functional integration (higher mean degree, higher clustering coefficient, and higher efficiency) for the sit-to-stand task (degree, *t-test: p <* 0.001, Fig. 5). On the other hand, linear and nonlinear muscle networks agree on inherent patterns of functional integration for the unilateral tasks (Fig. 3, 4).

This observation highlights the importance of both linear and nonlinear decoding of neural information sharing in the peripheral nervous system. In other words, the results suggest that when functional muscle networks are used as a biomarker of neuromusculoskeletal health/recovery, both linear and nonlinear measures should be studied concurrently to observe both linear and nonlinear synergistic patterns (which can be very different).

Capitalizing on this observation, in Fig. 6, a twodimensional functional network plane is proposed that includes both linear and nonlinear features. Each node in this plane represents a connectivity edge. As can be seen, the two-dimensional plane spans the considered task space, and the tasks form representative clusters in this 2D space. The bilateral clusters are readily separable in the 2D space.

As mentioned before, in this work, we also collect 64channel EEG data to observe if and how the muscle network behavior follows cortical involvement during the targetted functional tasks. The results can be seen in Fig 7. It should be noted that the higher nonlinear connectivity during the sit-to-stand task, when compared with the walking task, corresponds to the higher cortical activation (when comparing sit-to-stand with walking, Fig. 1c). This observation suggests that changes in the nonlinear connectivity mapping may better represent alterations in cortical activation during functional tasks, and this again highlights the importance of considering nonlinear measures of connectivity when evaluating information segregation and integration at the peripheral nervous system level.

## IV. Conclusion

In this paper, we compared linear versus nonlinear muscle networks for the dynamic lower-limb task. The results suggest that the two types of network scans would uncover different types of synergistic information sharing at the periphery. We proposed a two-dimensional connectivity plane composed of linear and nonlinear features and showed that it could span lower limb task space. Studying the concurrent EEG, it is shown that the activation map at the *δ* band may explain differences between the two core tasks, i.e., walking versus sit-to-stand. The study was limited by the number of subjects. The future line of research includes data collection and statistical analysis of the observed phenomena in this case study.

